# System-level health profiling from blood DNA methylation with explainable deep learning

**DOI:** 10.1101/2025.10.13.681975

**Authors:** David Martínez-Enguita, Thomas Hillerton, Julia Åkesson, Maria Lerm, Mika Gustafsson

## Abstract

Genome-scale DNA methylation (DNAm) profiles capture organismal physiology, but most predictive models lack transparency and multi-level applicability. Here we develop an explainable framework that quantifies respiratory, cardiovascular, and metabolic status as bounded health scores (0–1) derived from sex-specific clinical reference ranges and disease penalties, and then predicts these scores from whole-blood DNAm. Using Generation Scotland and case-control samples (n = 14,496 individuals), we screened 39 covariates for disease relevance and DNAm predictability, yielding system- relevant panels that were aggregated into scores. We compressed DNAm profiles with a protein-interaction-guided autoencoder, and trained health predictors on 128- dimensional embeddings using fully connected networks. On held-out samples, models reproduced the composite scores with strong rank agreement (Spearman ρ = 0.87, R^2^ = 0.71 for respiratory health; ρ = 0.82, R^2^ = 0.66 for cardiovascular; ρ = 0.81, R^2^ = 0.64 for metabolic) and recover expected population structure in a generally healthy cohort, with clear separation between “single-system low” and “multi-system low” phenotypes, and graded coupling across systems without redundancy. Further, the top features retrieved from each explainable predictor aligned with system biology: airway epithelial repair, hypoxia and inflammatory trafficking for respiratory; endothelial remodeling and cardiomyocyte programs for cardiovascular; glucose-lipid metabolism and metaflammation for metabolic. These results show that DNAm embeddings can yield accurate, transparent, and system-aware health profiling from blood, providing actionable summaries while revealing the molecular processes the models use to infer multi-system status. This approach positions DNAm embeddings plus interpretable penalty targets as a practical bridge from epigenomic signal to system-level triage and is extensible for evaluation in larger, more diverse cohorts.

## 1 INTRODUCTION

High-performance predictive models in medicine often face a major barrier to clinical translation: they can be accurate yet rely on poorly explained foundations that clinicians struggle to trust or act upon. In high-stakes domains like healthcare, understanding why an AI model makes a prediction can be as important as the prediction itself (Tonekaboni *et al.*, 2019) (Rao and Khan, 2022). Besides impeding acceptance, lack of interpretability raises safety and ethical concerns, since opaque models might latch onto spurious correlations or biases undetectable to users. Moreover, patients often present with multiple co-morbid conditions, with an estimated 42% of adults in the U.S. presenting at least two chronic diseases (Schiltz, 2022), underscoring that integrative models that can provide an actionable view of patient health rather than isolated predictions for single outcomes are needed. Thus, the development of transparent and interpretable models is necessary if they are to serve as reliable clinical decision support tools.

Efforts to address this challenge have given rise to various explainable modeling approaches in biomedical AI. A reasonable approach is to design inherently interpretable models that incorporate domain knowledge into model architecture, rather than relying solely on post-hoc explanations (Rudin, 2019). For example, simple logistic regression classifiers but also attention-based and case-based neural networks can at present determine relevant features or prototypical cases to justify their outputs, helping to understand model reasoning (Ghassemi, Oakden-Rayner and Beam, 2021). Within genomics, researchers are integrating biological priors to improve interpretability without sacrificing performance. For example, Kim et al. (2025) introduced an explainable deep learning framework called PROMINENT that predicts phenotypes from DNA methylation (DNAm) by integrating gene-level methylation features with known pathways. This approach improved accuracy over earlier deep models like MethylNet (Levy *et al.*, 2020) and, importantly, used pathway-level priors and SHAP value analyses to identify crucial genes and pathways involved in diseases like asthma and pulmonary fibrosis. Emerging work is now exploring multi-disease signatures (Zhang *et al.*, 2024) and multi-omics integration with explainable AI, demonstrating that transparent AI can reveal novel mechanisms. A recent study by Li et al. (2025) combined deep graph representation learning with explainability to uncover aging-related molecular networks using large single-cell and DNAm data to discover a ribosomal gene module and an inflammatory pathway driving aging. These advances illustrate the clear trend from clinical risk scores to multi-omics models that strive to be both accurate and interpretable.

In this study, we present a system-level health profiling approach using whole blood DNA methylation and explainable deep learning, aiming to enable a comprehensive health assessment. We define three quantitative health scores (respiratory, cardiovascular, and metabolic) as bounded indices (0 to 1) that summarize deviation from an optimal healthy state in each physiological system. Each score is constructed by aggregating multiple domain-specific clinical and phenotypic parameters into a single penalty-based metric. For example, the respiratory score integrates lung function measures and respiratory disease status, whereas the cardiovascular score incorporates blood pressure readings and cardiac disease history (with higher penalties for values outside healthy ranges). These system-level scores serve as targets for predictive modeling. To leverage the high-dimensional DNAm data, consisting of hundreds of thousands of CpG sites, in an efficient and biologically informed way, we first compress the methylation profiles into embeddings using a network-coherent autoencoder. This autoencoder is pre-trained on a large compendium of methylation data with a protein-protein interaction (PPI) network prior, so that the latent features are enriched for functionally independent and biologically meaningful patterns. On top of these embeddings, we train fully connected deep neural networks for each score to predict respiratory, cardiovascular, and metabolic health. The models are able to achieve strong accuracy (Spearman ρ = 0.81–0.87 for held-out test samples, R^2^ = 0.65– 0.71) in reproducing the composite health scores from DNAm alone. To ensure these DNAm-based predictors have learned relevant biological mechanisms, we apply a perturbation-based “light-up” analysis to yield an importance ranking of CpGs and their associated genes for each system’s prediction. The results show that our models capture system-specific biology: the top-activated methylation features for the respiratory predictor map to genes involved in lung function, hypoxia response, and airway inflammation; features for the cardiovascular predictor are enriched in pathways for vascular endothelial function, cardiac muscle development, and wound healing; and for the metabolic health predictor they relate to insulin signaling, lipid metabolism, and chronic inflammation. By linking model outputs to specific genes and pathways, our explainable health score predictors provide both a quantitative risk assessment and a window into the underlying molecular mechanisms. This system-level profiling framework, integrating DNAm with explainable deep learning, represents a novel step toward actionable AI in precision medicine, offering clinicians a tool that establishes multi-system health status from blood and allows to identify candidate mechanisms behind reduced respiratory, cardiovascular, or metabolic health, as reflected in their epigenomic profile.

## 2 MATERIALS AND METHODS

### 2.1 Data collection and preprocessing

Whole-blood DNA methylation samples (n = 18,857) from the Generation Scotland cohort (Smith *et al.*, 2006), generated on the Illumina Infinium MethylationEPIC BeadChip, were obtained as intensity data (IDAT) files, together with participant-level phenotypic and clinical data. Raw IDAT files were preprocessed in R. Beta values were derived with *minfi* (v1.54.1), after which between-sample technical variation at the intensity level was reduced using Gaussian mixture quantile normalization (GMQN) (Xiong *et al.*, 2022), followed by within-array beta-mixture quantile normalization (BMIQ) to correct Type II probe bias. Quality control and probe filtering were performed with *ChAMP* (v2.34.0) by removing (i) non-CpG probes, (ii) probes containing a single nucleotide polymorphism (SNP), (iii) probes mapping to the X or Y chromosome, (iv) multi-hit probes, and (v) probes not present on both Illumina 450K and EPIC arrays. Missing beta values were imputed by k-nearest neighbors (k = 10) using *bnstruct* (v1.0.15). From the initial cohort, we retained a complete-case analysis set of 11,867 samples with no missing values across 39 predefined phenotypic and clinical covariates plus ICD-coded disease information of interest. When applicable, covariates were numerically encoded and min-max scaled to [0, 1] prior to penalty calculations.

In addition, to enhance generalizability across cohorts, we curated publicly available case-control datasets from the Gene Expression Omnibus (GEO) repository. We screened case-control studies with human whole-blood DNA methylation profiles relevant to the respiratory, cardiovascular, or metabolic domains and identified 2,629 eligible profiles: n = 447 cardiovascular-related (e.g., atherosclerosis, coronary artery ectasia, stroke), n = 1,462 metabolic-related (e.g., obesity, type 1 and type 2 diabetes), and n = 720 respiratory-related (e.g., asthma, acute respiratory distress syndrome, severe COVID-19). These sample sets were integrated with the Generation Scotland complete-case analysis set, yielding a combined training cohort of 14,496 samples, each represented by 384,629 CpG probes.

### 2.2 Development of system-level health scores: respiratory, cardiovascular, and metabolic

We define a system-level health score as a bounded index *s*∈[0, 1] that quantifies an individual’s deviation from a reference healthy state for a given physiological system using a penalty-based framework. Considering the prespecified panel of domain- relevant phenotypic and clinical covariates *K_s_*, for each system-specific covariate *k* within system *s*∈{respiratory, cardiovascular, metabolic} for sample *i*, we assigned a monotonic penalty *p_sk_*(*x_ik_*) that is zero within a sex-dependent reference interval and increases with distance outside that range. ICD-coded disease contributed an additional system-specific penalty when applicable: zero if absent, and increasing monotonically with the severity of the condition. A composite system-level penalty *P_is_* can thus be calculated as the weighted sum of domain penalties *p_sk_*:

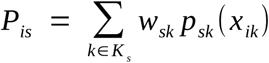

with the corresponding system-level health score S_is_ computed as the min-max reverse of *P_is_* across all training samples:

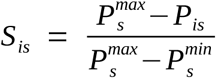

with larger composite penalties producing a lower health score (S_is_ ∼ 0), and lower or null penalties indicating better system-level health (S_is_ ∼ 1). We applied this approach across every health score, with differences only in the selection of the covariate panel, weights, and training sets. Reference ranges for covariates were defined according to guidelines from the World Health Organization (WHO) and the National Heart, Lung, and Blood Institute (NHLBI, NIH).

The selection of covariates for each system-level score was performed in the Generation Scotland cohort using a three-step procedure. Starting from the curated panel of 39 phenotypic and clinical covariates, we assembled system-specific covariate sets for the respiratory, cardiovascular, and metabolic scores. First, disease relevance was assessed by quantifying the association between the penalty value for each covariate across samples and the presence of the score-matched ICD-coded disease in the target system. Covariates were ranked based on their absolute Pearson correlation to the closest disease category (Lung Disease for the respiratory score, Heart Disease for the cardiovascular score, Diabetes for the metabolic score). Second, predictability from DNAm data was evaluated by training embedding-to-covariate deep neural network predictors using the 128-dimensional DNAm embeddings. Covariates with low performance were excluded. Third, to ensure interpretability we enforced non-overlap between score covariates, so that each retained covariate was used in at most one system.

### 2.3 Training of embedding-based health score predictors

We generated latent representations (embeddings) for all samples by compressing their DNAm profiles using a pre-trained, three-layered DNAm autoencoder with a 128- dimensional bottleneck. The autoencoder was trained on a large, pan-tissue compendium of DNAm profiles, as described in Martínez-Enguita *et al.* (2025), with its latent space constrained by a protein-protein interaction (PPI) graph constructed from high-confidence STRING v12 interactions (combined score ≥ 700) and RoseTTAFold2 interactions (probability ≥ 0.5), totaling 18,650 unique genes and 559,921 edges. We obtained embeddings for the combined training cohort, which served as fixed-length inputs to the downstream health score predictors.

System-specific predictors for respiratory, cardiovascular, and metabolic health scores were trained as fully-connected deep supervised neural networks (DNNs) of three hidden layers, with 128 hidden units each. Each dense layer was followed by batch normalization (momentum = 0.99, ε = 1e-3), leaky ReLU activation (α = 0.3), and dropout (rate = 0.10). Weights were initialized with He uniform, and dense layers were L2-regularized (λ = 1 x 10^-3^). The output layer used a sigmoid activation to constrain scores to [0, 1]. Models were trained to minimize mean squared error (MSE) with Adam (learning rate = 1 x 10⁻⁴, default β parameters) under an exponential decay schedule (decay rate = 0.95 every 10^3^ steps), using a batch size of 512 and early stopping with a patience of 1,000 epochs based on validation loss. Samples were randomly partitioned into training, validation, and test sets (64%, 16%, and 20%, respectively).

Performance on the test set was measured using the coefficient of determination (R^2^), mean absolute error (MAE), median absolute error (MedAE), root mean square error (RMSE), Pearson correlation coefficient (PCC, *r*), Spearman correlation coefficient (SCC, ρ), and margin of error for an individual prediction (measured as the 95% confidence interval), from *scikit-learn* v1.3.2 and *scipy* v1.10.1. All neural network models were implemented in Keras 2.4.3 running on TensorFlow 2.4.0 (TensorFlow- GPU 2.2.0) under Python 3.8.10.

### 2.4 Penalty optimization via cyclic coordinate ascent

To enable the application of the penalty approach to samples outside Generation Scotland (e.g., GEO datasets), we designed a penalty optimization scheme based on cyclic coordinate ascent (CCA). Because covariate metadata are incomplete, the appropriate penalties are unknown a priori. However, an exhaustive evaluation of all possible penalty combinations would be computationally prohibitive. Therefore, we followed a CCA-based approach since its coordinate-wise greedy updates can capture interaction effects pragmatically while keeping the evaluation burden minimal. First, we predefined discrete three penalty severity levels (severe, moderate, mild) for each covariate, scaled to the expected impact of that condition or disease subtype relative to the rest of included conditions. Then, we applied CCA to select the combination that maximized case-control separation in held-out Generation Scotland samples, measured as area under the receiver-operating curve (AUC), non-parametric effect size (Cliff’s delta) and Wilcoxon rank-sum P-value, in this order of priority. We iterated once over every condition and trained health score predictors on the selected target penalty combination. At every iteration, for condition *j*, we held all other penalties fixed and evaluated the alternative severity levels, replacing the current level with the one yielding the best test-set discrimination. We repeated the process until no additional improvement could be achieved. The final configuration reported is the one achieving the best disease-control separation by the established metrics.

### 2.5 Identification of top-activated genes from health score predictors using light- up analysis

The latent space of the trained health score neural networks was interrogated using “light-up” analyses (Dwivedi *et al.*, 2020) (Martínez-Enguita *et al.*, 2023) to obtain importance rankings of the input features (CpGs). The “light-up” technique is a perturbation-based interpretation method used to explain the internal representations learned by supervised DNNs. With model weights fixed, we systematically perturbed individual CpG sites in a representative DNAm profile and forward-propagated the changes through the pre-trained encoder and the system-specific predictor, recording the resulting change in the predicted health score (Δŷ) as a measure of feature importance. We evaluated two perturbations for each CpG *j* (hypermethylation, *β_j_* = 1; and hypomethylation, *β_j_* = 0) while holding all other probes constant, and then ranking CpGs by the magnitude of their effect on the corresponding health score prediction objective. The reference profile was defined as the medoid of the Generation Scotland cohort (the sample with minimal total Euclidean distance to all other samples). For each health score model (respiratory, cardiovascular, metabolic), we retained the top 1% of CpGs by relative importance and mapped them to their associated genes. We refer to these as the top-activated genes for the respective predictor, which were subsequently tested for functional enrichment analysis in relevant biological processes.

### 2.6 Gene annotation and functional enrichment analysis

Mapping of CpG probes to genes was performed with the Infinium MethylationEPIC v1.0 B4 Manifest File and probe annotation files provided by the *ChAMPdata* (v2.40.0) R package. Conversion and annotation of gene identifiers across Gene Symbols, Ensembl IDs, and NCBI IDs were carried out using *AnnotationDbi* (v1.66.0), *biomaRt* (v2.60.1), and *org.Hs.eg.db* (v3.19.1) R packages.

Functional enrichment analysis of top-activated genes from each health score prediction model in Gene Ontology terms of the Biological Process category (GO-BP) was conducted using the *clusterProfiler* R package (v4.12.6) with default parameters. Terms with and FDR-adjusted P < 0.05 were considered significantly enriched and were grouped into manually curated term categories by relevance to the respective health score.

### 2.7 Statistical analysis and figure editing

Statistical analyses and data processing were performed in R 4.5.1, within RStudio 2024.12.1, and Python 3.8.10. The threshold for significance corresponds to P-value < 0.05, unless otherwise stated. LOESS (locally estimated scatterplot smoothing) was used to fit nonparametric trends. Pairwise associations between system-level scores were quantified with Spearman’s rank correlation (ρ) and two-sided P-values.

Uncertainty was summarized with nonparametric bootstrap with 95% confidence intervals (10,000 resamples of paired observations) using the *boot* R package (v1.3-31).

Figures were created using *ggplot2* v3.5.2 R package, and *seaborn* (v0.11.1) and *matplotlib* (v3.4.2) Python libraries. Post-processing and panel layout adjustments for improved visualization were done in Inkscape v0.92.

## 3 RESULTS

To model multi-system health in a clinically meaningful way, we designed an end-to-end data-driven pipeline to derive composite scores that summarize respiratory, cardiovascular, and metabolic health status (Fig. 1). First, we screened 39 candidate phenotypic and clinical traits in the Generation Scotland cohort (n = 11,867, subset with complete trait data) for both disease relevance and predictability from whole-blood DNAm embeddings learned by a protein-interaction-guided autoencoder (Martínez- Enguita *et al.*, 2025). We converted the covariates selected for each system into penalties based on sex-adjusted healthy intervals, which were then aggregated into system-level health scores. Respiratory-linked traits chosen were spirometry test measurements: FEV (Forced Expiratory Volume, Pearson correlation of FEV penalty with lung disease = 0.169, Pearson correlation of true vs. predicted FEV from DNAm embeddings = 0.763), FVC (Forced Vital Capacity, cor_disease_ = 0.142, cor_prediction_ = 0.760), and FEF (Forced Expiratory Flow, cor_disease_ = 0.186, cor_prediction_ = 0.597). Cardiovascular covariates selected were average systolic (Pearson correlation of penalty with heart disease = 0.279, cor_prediction_ = 0.528) and diastolic blood pressure (cor_disease_ = 0.168, cor_prediction_ = 0.409). Metabolism-associated traits identified were waist-hip-ratio (whr, Pearson correlation of penalty with diabetes = 0.185, cor_prediction_ = 0.603) and body fat percentage (cor_disease_ = 0.193, cor_prediction_ = 0.691). As could be expected, age showed the strongest correlation with disease across covariates (Pearson correlation of age penalty with disease presence = 0.315) and was therefore included in all scores as a general indicator of health. Other covariates were not correlated with disease (cor_disease_ < 0.1) or could not be reliably determined from whole blood DNAm data (cor_prediction_ < 0.3) and were thus discarded. The presence of a system-relevant disease contributed an additional penalty depending on the severity of the condition. The scoring strategy was extended to relevant case-control GEO datasets (n = 2,629) by estimating the same penalties from DNAm alone, enabling the computation of scores in datasets lacking phenotype information. Using these health scores as supervised targets, we trained DNAm-based deep neural network predictors, yielding accurate and explainable models whose internal importance profiles recover biological processes aligned with each system’s pathophysiology.

**Figure 1.**
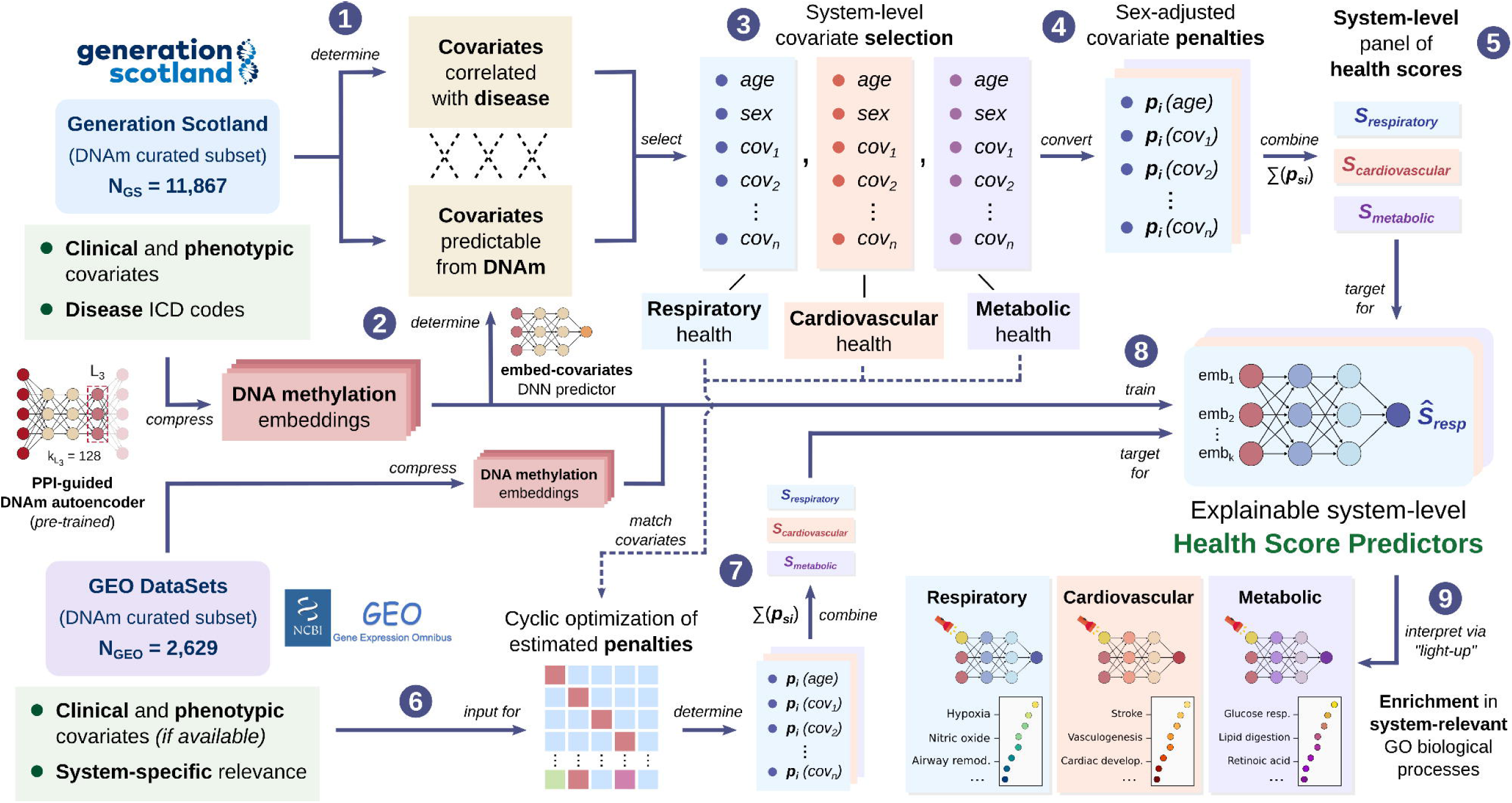
Graphical summary of the workflow for the training of explainable system-level health score predictors. Generation Scotland clinical and phenotypic covariates were analyzed for disease-association (1) and predictability from whole blood DNA methylation PPI-guided-autoencoder embeddings (2) to select system-specific representatives of respiratory, cardiovascular, and metabolic health (3). The selected covariates were converted into penalties using standard sex-dependent healthy ranges (4), which were then combined into system-level health scores (5). Publicly available samples from GEO without covariate information were assigned penalty estimates for the pre-selected system-specific covariates, which were optimized for disease relevance (6) and combined into health scores (7). Explainable supervised deep neural network models can thus be trained on DNA methylation embeddings to predict each health score (8). Lastly, the trained predictors can be interpreted using “light-up” to identify the biological processes captured in their learned representations (9).

### 3.1 System-level scores capture shared but distinct physiological dimensions and stratify selective and global impairment

Across Generation Scotland individuals, we observed that the three system-level scores captured a coherent portrait of physiological status. All scores span the full 0–1 range but show heavily right-shifted distributions, consistent with a generally healthy cohort (Fig. 2a**-c**). The respiratory score concentrates tightly near the upper end (median = 0.85, interquartile range [IQR] = 0.75–0.95) with few low values (only 6.9% of individuals below a score of 0.60), indicating limited impairment in the population and suggesting a ceiling region above 0.90 where very small decreases may still be clinically meaningful. Comparatively, the cardiovascular score distribution is more dispersed (median = 0.81 [0.68–0.92]), which is compatible with common subclinical variability in blood pressure and vascular risk and suggests the score can grade risk across a wide “gray zone” (between 0.50 and 0.80), not only flag extreme cases (2.0% of individuals below 0.4).

**Figure 2.**
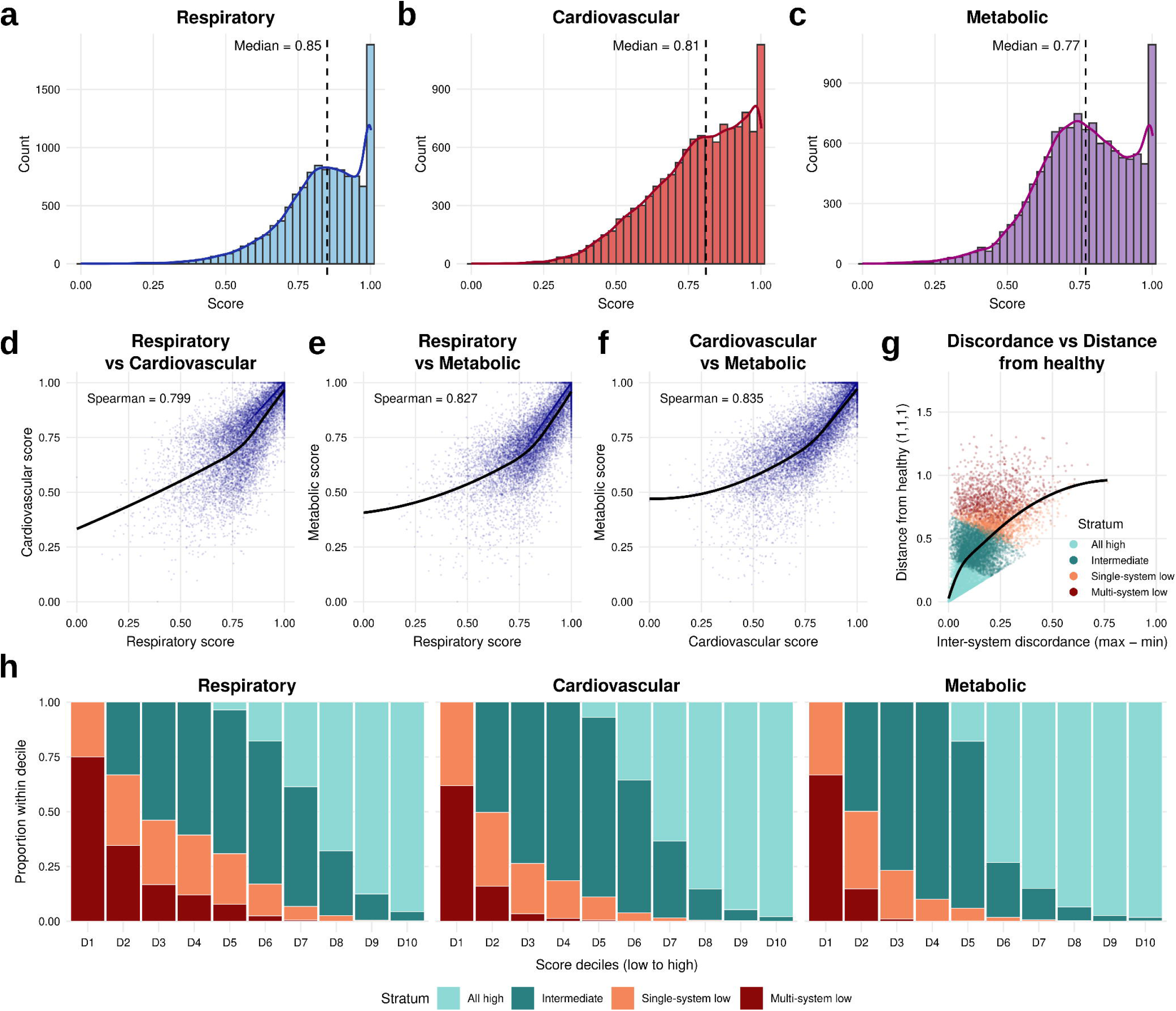
System-level health scores and cross-system dependencies. **(a-c)** Marginal distributions of the respiratory, cardiovascular, and metabolic health scores (0 to 1 scale, where 1 is the healthiest) on Generation Scotland individuals. Bars show sample counts, with the frequency outline overlayed. Dashed vertical lines mark the median across samples. **(d-f)** Pairwise scatterplots of the relationships between system scores. Points are individuals, black curves are LOESS trend lines. Spearman rank correlations (ρ) are reported inside panels. **(g)** Inter-system discordance vs. distance from health. The x-axis shows health score discordance, measured as the difference between the maximum and minimum health scores per sample. The y-axis shows Euclidean distance from the healthy corner (1,1,1). Points are colored by intrinsic health categories or strata (All high = light teal, Intermediate = teal, Single-system low = orange, Multi-system low = dark red) using pre-specified thresholds (high ≥ 0.80; low < 0.60). **(h)** Composition of strata across deciles of each score (D1 = lowest to D10 = highest). Bars represent the proportion of individuals per stratum within each decile.

Last, the metabolic score (median = 0.77 [0.66–0.89]) displays the longest left tail (14.7% of individuals below 0.6) with a visible shoulder at intermediate values (around 0.70–0.85, 33.1% of individuals), implying the existence of a sizable subgroup that departs from strictly healthy reference ranges. This could be expected given the background burden of obesity and dysglycemia in modern populations.

Pairwise relationships between scores showed broadly similar positive associations across systems (Fig. 2d**-f**). Individuals who were healthy in one system tended to be healthier in the others (Spearman ρ above 0.8 in all system pairs). However, the non- perfect correlations and residual variance at intermediate values indicate that each score contributes non-redundant, system-specific information. Cardio-metabolic coupling was the strongest (Spearman ρ = 0.84 [0.83–0.84]), while the respiratory-cardiovascular (ρ = 0.80 [95% CI 0.79–0.81]) and respiratory-metabolic (ρ = 0.83 [0.82– 0.83]) correlations were slightly lower but comparable. All were monotonic and statistically robust (P < 2.2 x 10^-6^ in all cases). The fitted LOESS curves suggest a mild non-linearity, since concordance is flatter near the healthy range and steepens once any score drops below ∼0.8, consistent with multi-system deviations emerging together as health declines. Notably, individuals with high respiratory values still show variability in cardiovascular or metabolic health. Among the subset with respiratory score ≥ 0.80 (n = 7,568, 63.8%), 25.8% had cardiovascular scores < 0.80 and 34.7% had metabolic scores < 0.80. Together, 40.6% had at least one of these below 0.80. Within this subgroup, cardiovascular and metabolic scores remained high (medians = 0.89 and 0.85, respectively; SD = 0.12 for both), with a cardio-metabolic discordance of 0.05 [0.02–0.10]. Their restricted-range correlations with respiratory scores decreased with respect to the full-range (ρ = 0.75 and 0.78), underscoring that preserved respiratory status tracks overall health at the upper range but does not guarantee a parallel cardiovascular or metabolic status.

To summarize overall impairment and cross-system imbalance, we computed two complementary quantities for each individual: (i) distance from the healthy vertex (1,1,1), which increases as all three scores depart from health; and (ii) inter-system discordance, the range (max–min) of the three scores, which increases when one system is much worse than the others. Using high and low thresholds of score ≥ 0.80 and < 0.60 to stratify individuals, the map of discordance versus distance (Fig. 2g) shows four clinically distinct behaviors. “All high” (n = 4,492, 37.8%) and “Intermediate” (n = 4,583, 38.6%) cases cluster near the origin (low to medium distance to health, average All high = 0.12 and Intermediate = 0.41; and discordance, 0.06 and 0.13, respectively), implying a preserved or only mildly compromised function across systems. “Single-system low” cases (n = 1,608, 13.6%), characterized by higher discordance (0.24) and moderate distance from health (0.62), reflect the selective failure of one system against relatively healthy others. Within this group, most involve cardiovascular (45.1%) or metabolic (40.3%) impairment . Individuals presenting this score pattern could benefit from a targeted evaluation of the outlying system rather than broad intervention. “Multi-system low” cases (n = 1,184, 10.0%) concentrate at high distance (0.83) with modest discordance (0.22). This phenotype is compatible with multi-morbidity, probably requiring coordinated and multi-system management. The overall positive association between discordance and distance (ρ = 0.68) indicates that large imbalances across systems rarely occur in otherwise healthy individuals, while the separation of these strata demonstrates that discordance between system scores can add information beyond severity by distinguishing selective from global impairments.

The stratified composition across score deciles per system (Fig. 2h) supports these patterns. Respiratory scores show the earliest decrease from the “All high” stratum by D5–D6, with a shift to Intermediate or Single-system low, indicating a mid-range fragility of the respiratory status. Cardiovascular scores sit in between, with the transition away from All high occurring mainly around D6–D7. Metabolic scores, however, present a more bimodal distribution, as the bottom deciles (D1–D3) are enriched for Single- and Multi-system low, but All high remains predominant through the mid-high deciles.

Clinically, this suggests respiratory health declines tend to appear sooner in the spectrum, while metabolic impairment concentrates among the lowest scorers and cardiovascular changes unfold more gradually.

### 3.2 DNAm embeddings accurately predict system-level health and generalize across cohorts

Supervised DNNs trained on DNAm embeddings reproduced the three system-level health scores with strong out-of-sample performance (Fig. 3a). Predictive strength was highest for the respiratory score (Spearman ρ true vs. observed = 0.874, R^2^ = 0.709), followed by cardiovascular (ρ = 0.820, R^2^ = 0.655) and metabolic (ρ = 0.813, R^2^ = 0.639). Residual spread was largest at the lowest observed scores, a pattern expected when labels are derived from composite penalties that amplify small covariate errors. Importantly, the monotonic rank agreement (ρ ≥ 0.81 in all cases) indicates that the models can prioritize samples even when absolute calibration may require cohort-specific tuning.

**Figure 3.**
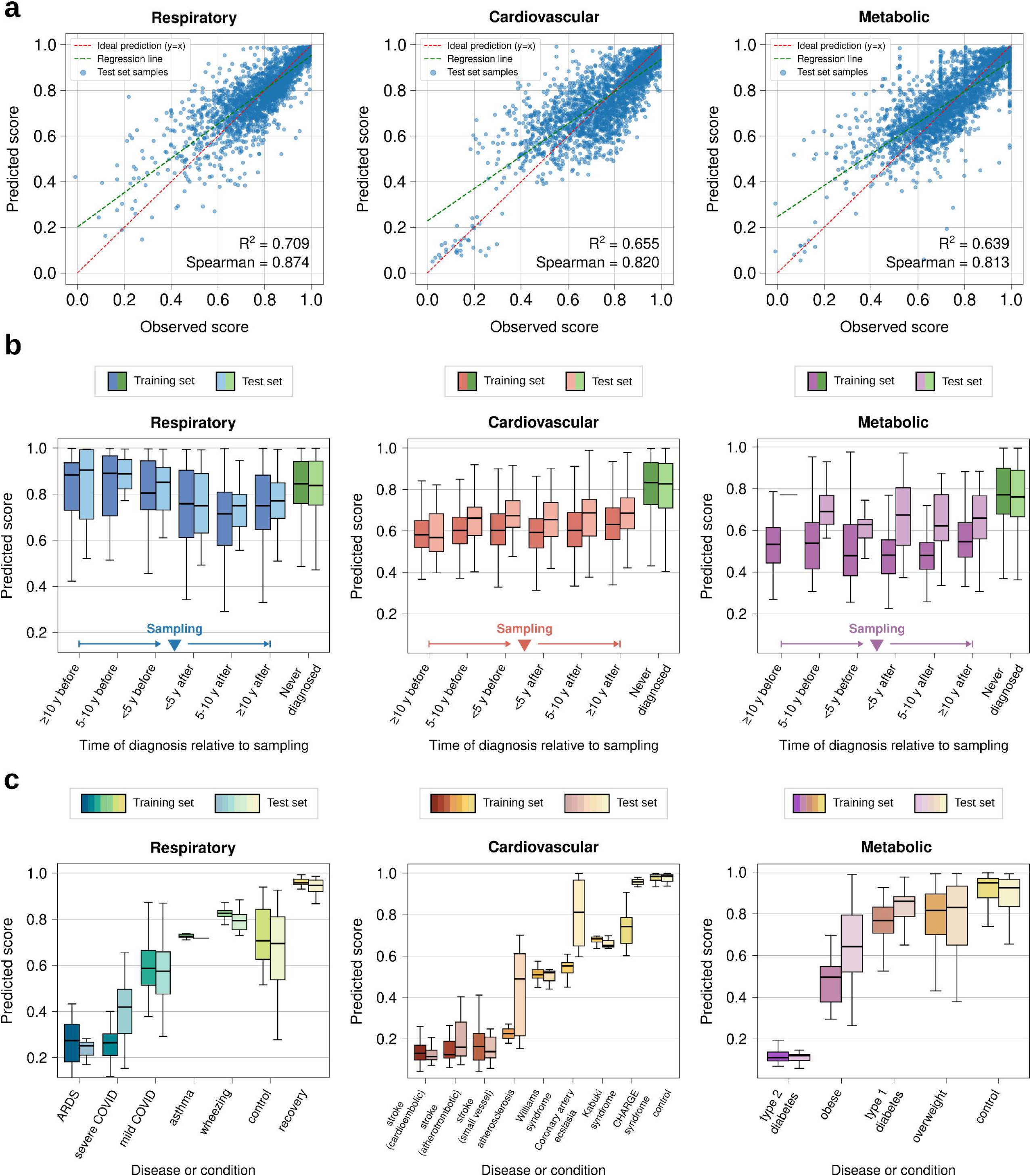
Performance of health score predictor models. **(a)** Observed vs. predicted health scores for system-level predictors (left to right: respiratory, cardiovascular, and metabolic) and performance on the test set, measured as explained variance (coefficient of determination, R^2^) and Spearman correlation. **(b)** Predicted health scores for Generation Scotland individuals with at least one system-relevant disease diagnosis, categorized by time of diagnosis relative to sampling, measured in intervals of five years before or after the sample was taken. Never-diagnosed individuals (controls) for system-relevant diseases are shown on the right side of every panel. **(c)** Predicted health scores for publicly available DNAm dataset samples from individuals with system-relevant diseases or conditions, categorized by decreasing order of severity (or increasing predicted health score). Darker-colored boxes correspond to training samples, light-colored boxes correspond to test samples.

For further fine-grained analysis, we grouped Generation Scotland participants with a system-associated ICD diagnosis (n = 3,071, 25.9%; divided into respiratory-associated diseases, n = 685, 5.8%; cardiovascular, n = 1,971, 16.6%; metabolic, n = 415, 3.5%) by the time of diagnosis relative to blood draw for DNAm sampling (Fig. 3b). The predicted scores tracked the clinical timeline in the expected direction. For each system, individuals diagnosed close to the sampling date or thereafter (“<5y after”) had the lowest median predicted health, whereas those diagnosed many years before sampling or never diagnosed showed progressively higher scores. This trend was visible in both training and held-out test subsets and was most pronounced for the cardiovascular and metabolic predictors, consistent with their broader dispersion in the population. The temporal gradient suggests that the models are sensitive to subclinical physiology that precedes formal diagnosis, raising the possibility of earlier risk flagging. Conversely, high scores among remote-diagnosis groups are compatible with disease resolution or effective management.

Applying the predictors to GEO datasets showed a general replication of the ordering of conditions by relative clinical severity established during the estimation of penalties (Fig. 3c). For the respiratory score, acute and severe phenotypes (e.g., acute respiratory distress syndrome, ARDS, and severe COVID-19) had the lowest predicted health (median = 0.25 [0.21–0.27] and 0.42 [0.30–0.50], respectively), intermediate chronic states (mild COVID-19, asthma, wheeze) scored higher (0.58 [0.48–0.66], 0.72, and 0.79 [0.75–0.82]), and samples without diagnosis were also moderately high but variable (0.69 [0.54–0.81]). Cardiovascular datasets showed a graded pattern ranging from severe events, like the different stroke types (medians between 0.11 [0.10–0.15] and 0.16 [0.12–0.28]), to medically managed states and finally to mainly young controls (0.99 [0.96–0.99]), again with clear separation between the diseased and non-diseased groups. The higher-than-expected scores for atherosclerosis (0.49 [0.21–0.61]) and coronary artery ectasia (0.81 [0.65–0.97]) may reflect treatment effects shifting DNAm- derived signals toward healthier values. For the metabolic predictor, the score distribution was consistent with differing pathophysiology and treatment. Type 2 diabetes ranked lowest (0.12 [0.10–0.13]), obesity (0.64 [0.52–0.80]) and type 1 diabetes (0.86 [0.79–0.88]) intermediate, overweight higher (0.83 [0.65–0.93]), and controls highest (0.93 [0.83–0.97]), in both training and test sets. These results support the validity and generalizability of the system-level DNAm-based health scores for comparative phenotyping and cohort categorization.

### 3.3 Explainable health-score predictors capture system-relevant biological processes

We interrogated each trained predictor with a perturbation-based “light-up” analysis that quantifies how directed changes at individual CpGs propagate to the model output (Methods). For each system, we ranked CpGs by importance, retained the top 1% as model-relevant features, mapped them to genes, and tested the resulting sets for Gene Ontology (GO) Biological Process enrichment (FDR-adjusted P < 0.05). The enriched terms (Fig. 4) provide a pathway-level readout of what biology the models use when estimating system-level health from whole blood DNAm embeddings.

**Figure 4.**
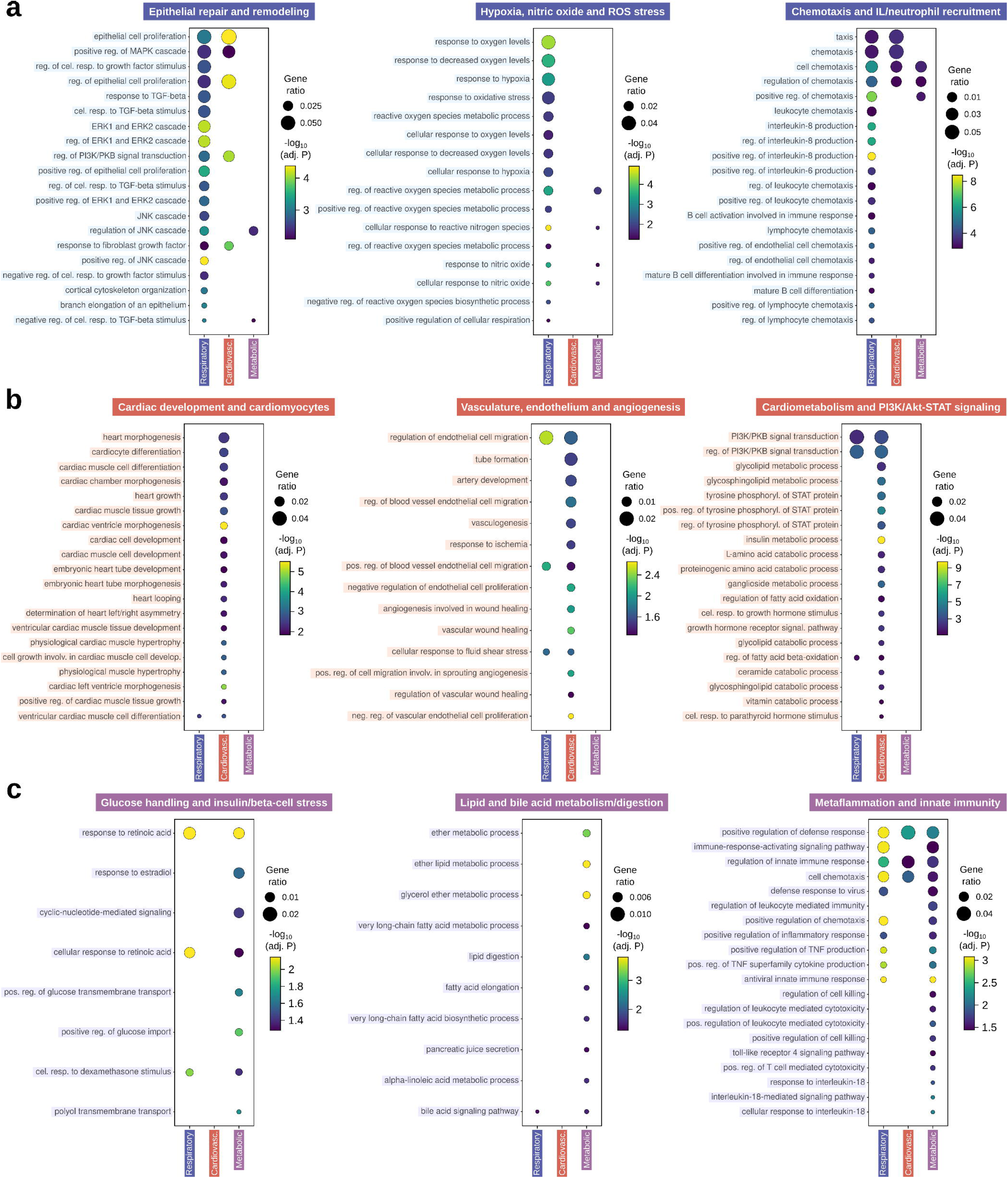
Enrichment in system-relevant GO biological process categories for DNA methylation signatures from interpretable health score predictors. Top significantly enriched (FDR adjusted P-value < 0.05) Gene Ontology (GO) terms of the Biological Process (BP) category for the gene signatures retrieved from the trained system-level health score predictors, grouped by relevant function. **(a)** Respiratory score predictor signature enrichment in GO-BP terms grouped into “Epithelial repair and remodeling”, “Hypoxia, nitric oxide and reactive oxygen species (ROS) stress”, and “Chemotaxis and interleukin/neutrophil recruitment”. **(b)** Cardiovascular score predictor signature enrichment in GO-BP terms grouped into “Cardiac development and cardiomyocytes”, “Vasculature, endothelium and angiogenesis”, and “Cardiometabolism and PI3K/Akt-STAT signaling”. **(c)** Metabolic score predictor signature enrichment in GO-BP terms grouped into “Glucose handling and insulin/beta-cell stress”, “Lipid and bile acid metabolism and digestion”, and “Metaflammation and innate immunity”. Dot size shows gene ratio; dot color shows enrichment significance, measured as negative log_10_ FDR-adjusted P-value. GO-BP terms found to be significantly enriched across other health score signatures are shown besides the main health score enrichment in each dot plot.

About the respiratory predictor, we observed it highlights programs central to airway integrity and regeneration (Fig. 4a), consistent with a sensitivity to obstructive and interstitial lung pathology. Enriched terms include the positive regulation of epithelial cell proliferation (FDR-adjusted P = 3.2 x 10^-4^), ERK/MAPK cascade (P = 8.6 x 10^-5^), and TGF-beta signaling (P = 7.0 x 10^-4^). These processes govern barrier maintenance, mucus-producing cell turnover and fibrotic remodeling, suggesting that the model leverages DNAm variation aligned with epithelial repair dynamics (present also in the cardiovascular model signature) in a biologically coherent manner. Other highly specific enriched terms such as responses to hypoxia (P = 6.3 x 10^-4^), to oxygen levels (P = 5.8 x 10^--5^), to nitric oxide (P = 4.5 x 10^-4^), and to oxidative stress (P = 9.5 x 10^-3^) indicate that the model captures vascular-alveolar gas-exchange stress and redox homeostasis, while nitric-oxide biology also modulates airway tone. Enrichment for lymphocyte chemotaxis (P = 2.2 x 10^-5^), B cell differentiation (P = 2.9 x 10^-4^), and interleukin-6 and interleukin-8 production (P = 1.5 x 10^-4^ and 3.2 x 10^-9^, respectively) may reflect airway inflammation and innate immune trafficking.

For the cardiovascular model (Fig. 4b), terms spanning cardiac muscle cell differentiation (P = 3.0 x 10^-3^) and development (P = 8.7 x 10^-3^), cardiac ventricle morphogenesis (P = 4.0 x 10^-6^), and cardiac hypertrophy (P = 1.2 x 10^-3^) point to lineage and structural programs of the myocardium. The gene signature retrieved is also strongly enriched for regulation of endothelial cell proliferation (P = 2.7 x 10^-3^), blood- vessel morphogenesis (P = 2.7 x 10^-2^), angiogenesis (P = 1.0 x 10^-2^), and response to ischemia (P = 2.9 x 10^-2^). These GO-BP terms hint towards the modeling of processes such as atherosclerosis, vascular remodeling, and plaque instability. Likewise, enrichments for PI3K/Akt and JAK/STAT signaling (P = 2.7 x 10^-5^ and 3.2 x 10^-6^), insulin-related metabolic processes (P = 4.2 x 10^-10^) and cellular responses to growth hormone (P = 3.6 x 10^-4^) suggest the model may capture the cross-talk between inflammatory signaling and cardiometabolic state that drives ventricular remodeling and vascular dysfunction. Given the clinical intersection of insulin resistance, inflammation and cardiovascular risk, the dependence on these pathways supports the model’s mechanistic plausibility and its sensitivity across the cardiometabolic spectrum.

Last, the metabolic health score predictor (Fig. 4c) was found to prioritize processes such as regulation of glucose transport (P = 1.3 x 10^-2^), response to retinoic acid (P = 8.0 x 10^-3^), and cyclic-nucleotide-mediated signaling (P = 3.5 x 10^-2^), potentially indicating the tracking of glycemic control and beta-cell workload. Similarly, enrichment for fatty-acid metabolic process (P = 2.0 x 10^-2^), very-long-chain fatty-acid biosynthesis (P = 2.0 x 10^-2^), lipid digestion (P = 4.3 x 10^-3^), bile-acid signaling (P = 2.4 x 10^-2^) and pancreatic juice secretion (P = 3.8 x 10^-2^) reflects hepatic and intestinal lipid handling, which are a hallmark of dyslipidemia and conditions such as non-alcoholic fatty liver disease (NAFLD). Other significantly enriched terms include positive regulation of defense response (P = 7.3 x 10^-3^), antiviral immune response (P = 8.9 x 10^-4^), leukocyte- mediated cytotoxicity (P = 1.3 x 10^-2^), TNF production (P = 7.3 x 10^-3^), and cellular response to interleukin-18 (P = 7.3 x 10^-3^). These immune-associated terms, partially present in the signatures extracted from the respiratory and cardiovascular predictors, point to a possible capture of the low-grade innate immune activation (“metaflammation”) that accompanies adiposity, insulin resistance, and fatty-liver disease.

Overall, the derived GO programs are biologically coherent with each system and provide interpretable anchors for the predictors. The score-specific pathway signatures show that the three predictors functionalize the PPI-guided DNAm embeddings to rely on biologically coherent processes: airway epithelial repair and inflammatory traffic for respiratory health, endothelial remodeling and cardiomyocyte programs for cardiovascular health, and glucose-lipid metabolism with metaflammation for metabolic health. These pathway-level insights support face validity and suggest concrete axes along which therapeutic intervention may shift the predicted health scores.

## 4 DISCUSSION

Our system-level health scores are designed to summarize respiratory, cardiovascular, and metabolic status by converting clinically interpretable covariates into monotonic penalties that increase with distance from sex-adjusted healthy intervals. This design makes two properties explicit: (i) every selected covariate retains its clinical meaning, and (ii) departure from health for any covariate lowers the score proportionally and cumulatively across traits. The three systems were chosen because they represent prevalent and interacting axes of morbidity with shared environmental and behavioral factors, such as smoking, adiposity, or inflammation, but distinct pathophysiology. Their relationships in the studied cohorts were found to be strong but non-identical, with a clear separation of “single-system low” versus “multi-system low” profiles. These patterns imply that the scores capture complementary biological signals and, when used together, have the potential to resolve phenotypes that a single score would miss. For example, metabolically-driven cardiometabolic risk despite preserved respiratory status, or isolated airway vulnerability in otherwise low-risk profiles. Practically, this supports a multi-signal decision strategy in which interventions are prioritized to the health score that first diverges, while continued surveillance focuses on the paired system most likely to decline next. Furthermore, age, the most universally disease-associated covariate, is incorporated into each score to reflect its broad influence on organ function and complement the system-specific variation captured by the remaining traits.

Across clinical timelines, predicted scores aligned with diagnostic proximity to the sampling, with lower values among individuals diagnosed near or after this timepoint and higher in those with remote or no diagnosis. This gradient suggests a certain level of sensitivity to subclinical physiology could be captured, and thus there is potential for early risk flagging and monitoring. Crucially, our predictors are not opaque: the examination of their inner activations exposed the sets of CpGs and associated genes that each trained health score model relies on, which in turn converged on pathway programs coherent with the corresponding system. In this way, the respiratory model emphasized epithelial repair, hypoxia and redox responses, and leukocyte trafficking, suggesting a capacity to detect systemic footprints of impaired oxygen delivery and redox imbalance in blood DNAm. Because neutrophil-dominant inflammation and the interleukin-axis activity typify chronic bronchitis and infectious exacerbations, the respiratory predictor’s reliance on these signals supports its physiological specificity to inflammatory airway disease. By contrast, the cardiovascular model prioritized endothelial remodeling, extracellular matrix and cardiac developmental and conduction programs. These DNAm signals likely index systemic regulation (e.g., endocrine or developmental pathways) or genetic and epigenetic predisposition that can affect myocardial structure and rhythm, which are consistent with the model’s ability to track variation across hypertrophy, conduction disease and heart-failure phenotypes. Their presence also indicates that signals of endothelial activation and repair are captured, aligning with the performance observed for ischemic and cerebrovascular disease. In turn, the metabolic model highlighted glucose responses and lipid handling alongside low-grade inflammatory signaling. These mechanisms, which are expected to shift across prediabetes and type 2 diabetes, match the predicted lower scores in diabetic cohorts. The reliance of the model on them aligns with a blood-based readout of systemic metabolic stress. In addition, the prominence of bile acid pathways further suggests a sensitivity to enterohepatic signaling that could be relevant to insulin resistance and obesity. We believe the processes identified provide robust mechanistic anchors for the model outputs, enabling hypothesis generation (e.g., whether interventions that modulate endothelial activation or oxidative stress might shift predicted scores) and enhancing clinical interpretability by mapping methylation-derived signals to recognizable function.

As limiting factors for this work, we first recognize that broader representation (across ancestries, age ranges, exposures, and underrepresented clinical phenotypes) is needed to further refine penalties and improve generalizability. Second, while performance is strong for a first-generation framework, richer priors and further calibration in prospective cohorts would be beneficial to increase the robustness of the score estimations. Third, the penalty calculation includes chronological age, but future iterations should evaluate biologically derived aging measures, such as DNAm-based epigenetic age, to separate calendar time from physiological decline. Methodologically, the approach is extensible: additional systems (e.g., renal, hepatic, neurocognitive or immune) can be incorporated by curating system-relevant covariates and penalties, and additional information layers can be integrated in the form of multi-omics or longitudinal constraints. Last, validation in larger and medically verified cohorts is required for the pipeline to be applied to complement routine labs, with system-specific thresholds tailored to triage and follow-up.

In sum, by coupling biologically guided targets with explainable DNAm-based predictors for respiratory, cardiovascular, and metabolic health, our framework delivers system- level, clinically readable health summaries from blood, an actionable step toward trustworthy epigenomic decision support in precision medicine.

## ACKNOWLEDGMENTS

Computational resources were granted by the National Academic Infrastructure for Supercomputing in Sweden (NAISS), including National Supercomputer Centre resources Berzelius (Berzelius-2024-5), Sigma (NAISS 2023/5-303), and Tetralith (NAISS 2024/5-385); and AIDA Data Hub science platform, part of SciLifeLab Bioinformatics Platform (National Bioinformatics Structure Sweden, NBIS).

## AUTHOR CONTRIBUTIONS

David Martínez-Enguita (Conceptualization [lead], Data Curation [lead], Formal Analysis [lead], Investigation [lead], Methodology [lead], Software [lead], Validation [lead], Visualization [lead], Writing – Original Draft [lead], Writing – Review & Editing [lead]), Thomas Hillerton (Formal Analysis [supporting], Investigation [supporting], Validation [supporting], Writing – Review & Editing [supporting]), Julia Åkesson (Formal Analysis [supporting], Investigation [supporting], Validation [supporting], Writing – Review & Editing [supporting]), Maria Lerm (Conceptualization [equal], Project Administration [equal], Resources [equal], Supervision [equal], Writing – Review & Editing [supporting]), Mika Gustafsson (Conceptualization [lead], Funding Acquisition [lead], Methodology [supporting], Project Administration [lead], Resources [lead], Supervision [lead], Writing – Review & Editing [supporting]).

